# Genetic diversity increases food-web persistence in the face of climate warming

**DOI:** 10.1101/2020.06.23.167387

**Authors:** Matthew A. Barbour, Daniel J. Kliebenstein, Jordi Bascompte

## Abstract

Genetic diversity provides the raw material for species to adapt and persist in the face of climate change. Yet, the extent to which these genetic effects scale at the level of ecological communities remains unclear. Here we experimentally test the effect of plant genetic diversity on the persistence of an insect food web under a current and future warming scenario. We found that plant genetic diversity increased food-web persistence by increasing the intrinsic growth rates of species across multiple trophic levels. This positive effect was robust to a 3°C warming scenario and resulted from allelic variation at two genes that control the biosynthesis of chemical defenses. Our results suggest that the ongoing loss of genetic diversity may undermine the persistence and functioning of ecosystems in a changing world.

**One Sentence Summary:** The loss of genetic diversity accelerates the extinction of inter-connected species from an experimental food web.

## Main Text

Gene-to-ecosystem processes sustain life on Earth. For example, genes encode information that determines an organism’s phenotype and fitness in the environment (*1*), which in turn plays a fundamental role in determining its trophic interactions with other species (*2, 3*). Similarly, the strength and organization of trophic interactions in a food web play an important role in maintaining species diversity across multiple trophic levels (*4, 5*). Predicting how living systems will respond to ongoing climate change, therefore, requires a mechanistic understanding of linkages across biological scales (*6, 7*). Given that the current rate of population extinction, and subsequent loss of genetic diversity, is orders of magnitude higher than the rate of species extinction (*8*), there is a pressing need to know to what degree this loss of genetic diversity will undermine the persistence of food webs in a changing world.

To understand how gene-to-ecosystem processes will respond to climate change, we used an experimental food web consisting of a plant (*Arabidopsis thaliana*), two species of aphids (*Brevicoryne brassicae* and *Lipaphis erysimi*), and a parasitoid wasp (*Diaeretiella rapae*) (Fig. 1A). These species form a naturally occurring food web (*10, 11*), which contains three of the most common interaction structures found in natural food webs, including resource competition (aphid-plant-aphid), apparent competition (aphid-parasitoid-aphid), and a food chain (plantaphid-parasitoid) (*12*). Interactions in this food web are partly mediated by a group of specialized metabolites called aliphatic glucosinolates (*10*). Extensive knowledge of genotype-to-phenotype causality in *Arabidopsis* aliphatic glucosinolates (Fig. 1B) provides a system to test how genetic change influences ecological interactions in a food web. We used 3 transgenic lines that recreate natural null alleles in the aliphatic glucosinolate pathway (*MAM1, AOP2*, and *GSOH*), which determine natural variation in aliphatic glucosinolates across multiple Brassicales species (*13–17*), in a common genetic background (Col-0 accession, Fig. 1B). Together, the Col-0 accession and transgenic lines reproduce most of the natural variation in the chemical phenotypes of *Arabidopsis* accessions that co-occur throughout Europe and determine fitness under field conditions (*1, 9, 18*). We created experimental plant populations from all possible combinations of *Arabidopsis* genotypes (*n* = 11 combinations across 60 experimental food webs), including monocultures (*n* = 4), two-genotype mixtures (*n* = 6), and the four-genotype mixture (*n* = 1). This experimental design allowed us to test both a general effect of genetic diversity and isolate the contribution of allelic variation at specific genes.

**Fig. 1.**
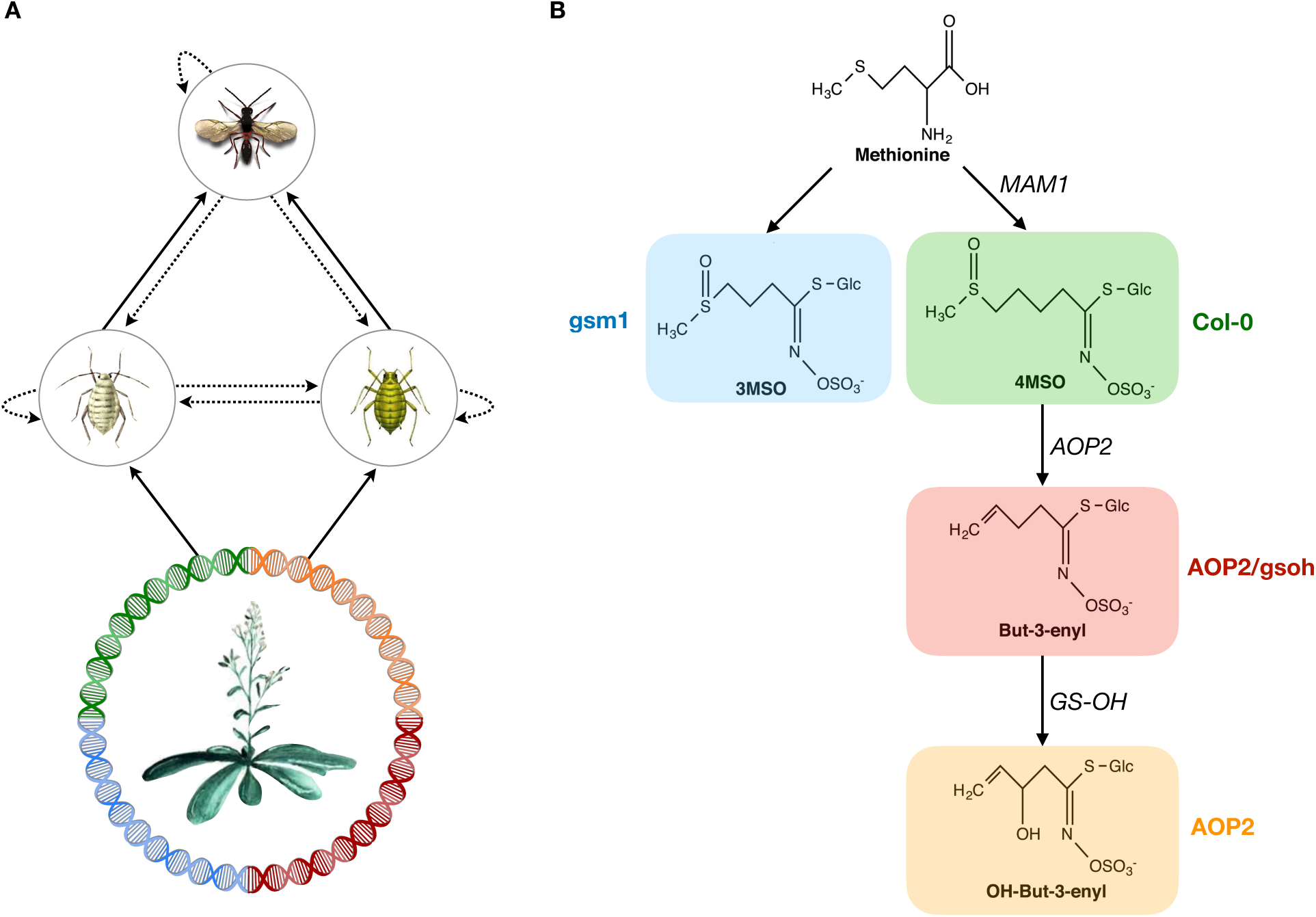
Study system. (**A**) In our experimental food web, the parasitoid *Diaeretiella rapae* (top) parasitizes the aphids *Brevicoryne brassicae* (left) and *Lipaphis erysimi* (right), and these aphids compete for their shared resource *Arabidopsis thaliana* (bottom). This food-web diagram represents our prior expectation of food-web structure, where solid and dashed arrows represent positive and negative effects, respectively. (**B**) To manipulate genetic diversity, we used 3 transgenic lines of *Arabidopsis* (gsm1, AOP2, AOP2/gsoh) that alter the biosynthesis of aliphatic glucosinolates in a common genetic background (Col-0 accession). The chemical phenotype (3MSO, 4MSO, But-3-enyl, or OH-But-3-enyl) of each *Arabidopsis* genotype depends on which genes (*MAM1, AOP2*, and *GS-OH*) have functional alleles (details given in note *9*). The glucosinolate pathway is adapted from Fig. 1 in ref. *1*.

To test the sensitivity of these genetic effects to climate change, we conducted the experiment under two different, constant temperature regimes (20°C and 23°C). We chose these temperatures to reflect the warming this food web is expected to experience over the next 25–50 years (*19*). More generally, warming can simultaneously modify processes at different biological scales (*6*). This is because temperature fundamentally determines an organism’s metabolic rate, which can scale up to alter the strength and stability of trophic interactions with other species (*20*).

After adding the experimental food web to each plant population, we tracked the population dynamics of each species for 17 weeks, allowing for multiple generations of aphids (∼16 generations) and parasitoids (∼8 generations). This allowed us to track critical transitions in the food web over time (Fig. 2). In the context of this paper, critical transitions refer to local extinctions that simplify the food web (*21*).

**Fig. 2.**
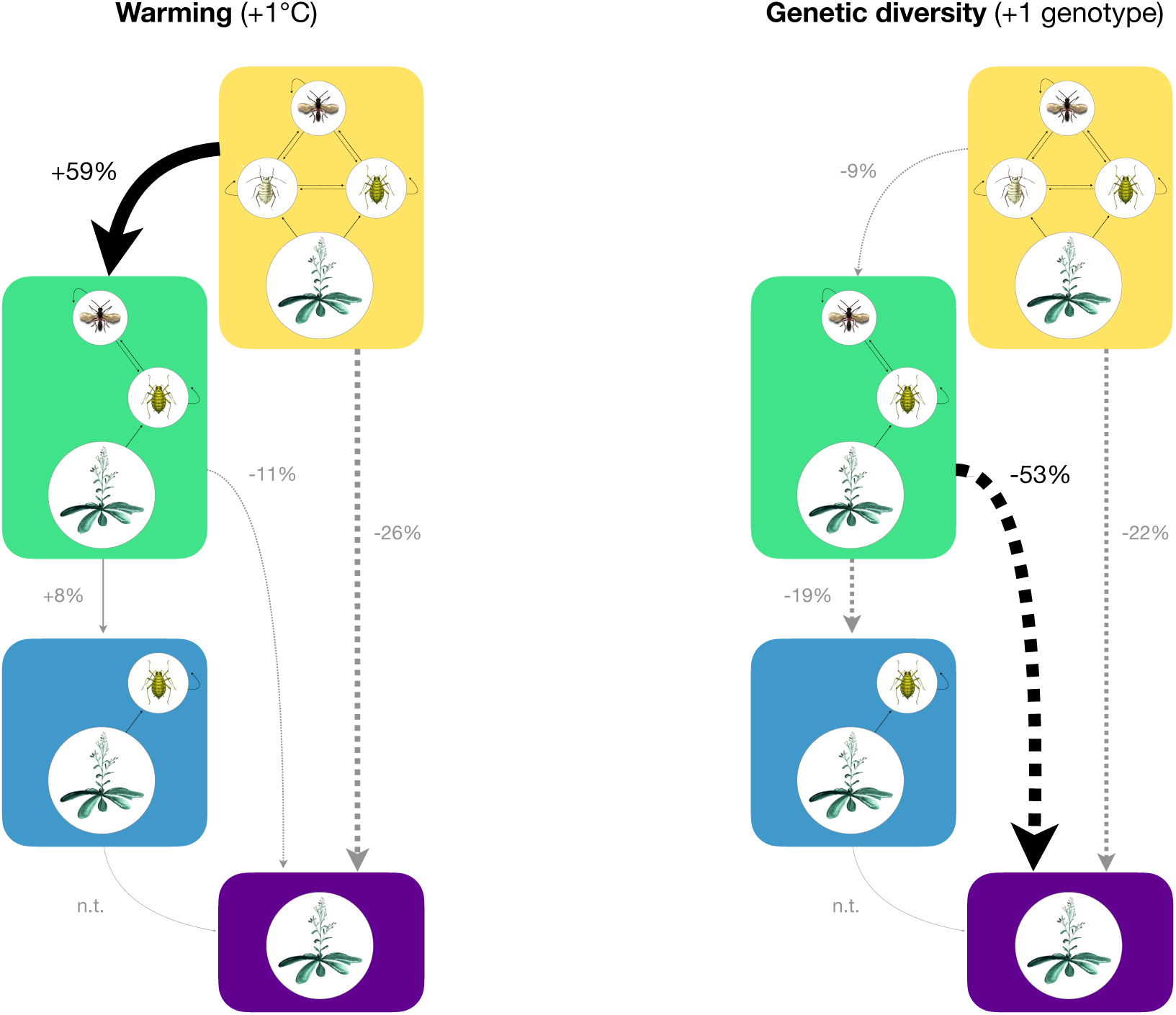
Critical transitions in the experimental food web. Each color corresponds to a different food-web structure, and arrows between two colors indicate a critical transition. The thickness of an arrow is proportional to the percent change in probability of a critical transition for every one unit increase in temperature (left) or genetic diversity (right). Solid arrows indicate positive changes, while dashed arrows indicate negative changes. Black arrows denote statistically clear effects (*P <* 0.05), while grey arrows are unclear. Rare critical transitions were not tested (labeled n.t.) for differences between treatments. Details on the statistical tests for each critical transition are provided in Tables S1–S2.

Warming had an immediate effect on food-web persistence (Fig. 2, Table S1). Specifically, a 1°C increase in warming accelerated the extinction of the aphid *Brevicoryne brassicae*, resulting in a simpler food chain (yellow to green in Fig. 2, S1–S3). However, warming did not alter the risk of a critical transition in the remaining food chain (green to either blue or purple in Fig. 2, Table S2). Although counterintuitive, warming increased the odds that the food web was reduced to only the aphid *Lipaphis erysimi* (blue in Fig. 2), rather than completely collapsing (purple in Fig. 2), by the end of the experiment (multinomial GLM ANOVA: *F*_2,20_ = 4.03, *P* = 0.034, Fig. S4).

Although warming accelerated the transition of the intial food web toward a simpler food chain, it was genetic diversity that governed the risk of a critical transition in the remaining food chain (Fig. 2, Table S2). Specifically, adding another genotype to the plant population reduced the risk of the food chain completely collapsing by 53% (green to purple in Fig. 2). This reduced risk of a critical transition tripled the odds of the food chain persisting to the end of the experiment (relative to complete collapse, multinomial GLM ANOVA: *F*_2,18_ = 3.61, *P* = 0.048, Fig. S4). Therefore, despite the initial effect of warming, the net effect of genetic diversity on food-web persistence was larger than that of warming (Fig. S4). Moreover, there was no clear evidence that temperature modified the effects of genetic diversity on any of the possible critical transitions (all *P* > 0.40 for ‘temp:rich’ in Tables S1–S2), suggesting that the positive effect of genetic diversity was robust to experimental warming.

To understand the source of this genetic diversity effect, we isolated the role of each specific genotype (Fig. 3). To do this, we leveraged the fact that the genetic diversity effect (adding an additional genotype in Fig. 3A) corresponds to the average genotype-specific effect (i.e., average of green, blue, red, and orange points in Fig. 3B). We focused on the effect of genetic diversity on the overall risk of a critical transition in the food chain (green in Fig. 3A), i.e., a transition to either an aphid only (blue in Fig. 3A) or completely collapsed state (purple in Fig. 3A). The rationale for this choice is that there was more certainty in this overall effect (*P* = 0.015) compared to the reduced risk of a complete collapse (*P* = 0.044 for green to purple transition in Fig. 2). We found that adding Col-0 or gsm1 to the plant population reduced the overall risk of a critical transition in this food chain by 45% and 48%, respectively (Fig. 3B). These genotypes differ in that Col-0 has a functional allele at the *MAM1* gene, whereas gsm1 has a non-functional allele. The *MAM1* gene controls the elongation of the glucosinolate side chain (Fig. 1B, ref. *13, 15*), and influences herbivory from aphids (*10*) and other insects (*15*). These independent contributions suggest that these genotypes have complimentary effects on food-chain persistence. Interestingly, Col-0 and gsm1 both lack a functional *AOP2* gene, which modifies the glucosinolate side chain (Fig. 1B, ref. *14, 16*). The two genotypes that have a functional *AOP2* gene had unclear effects on food-chain persistence (Fig. 3B). Analagous to a keystone species that determines species diversity in a food web (*22*), our results indicate that *AOP2* functions as a keystone gene in this food web by shaping species diversity through differences in food-web persistence (*23*).

**Fig. 3.**
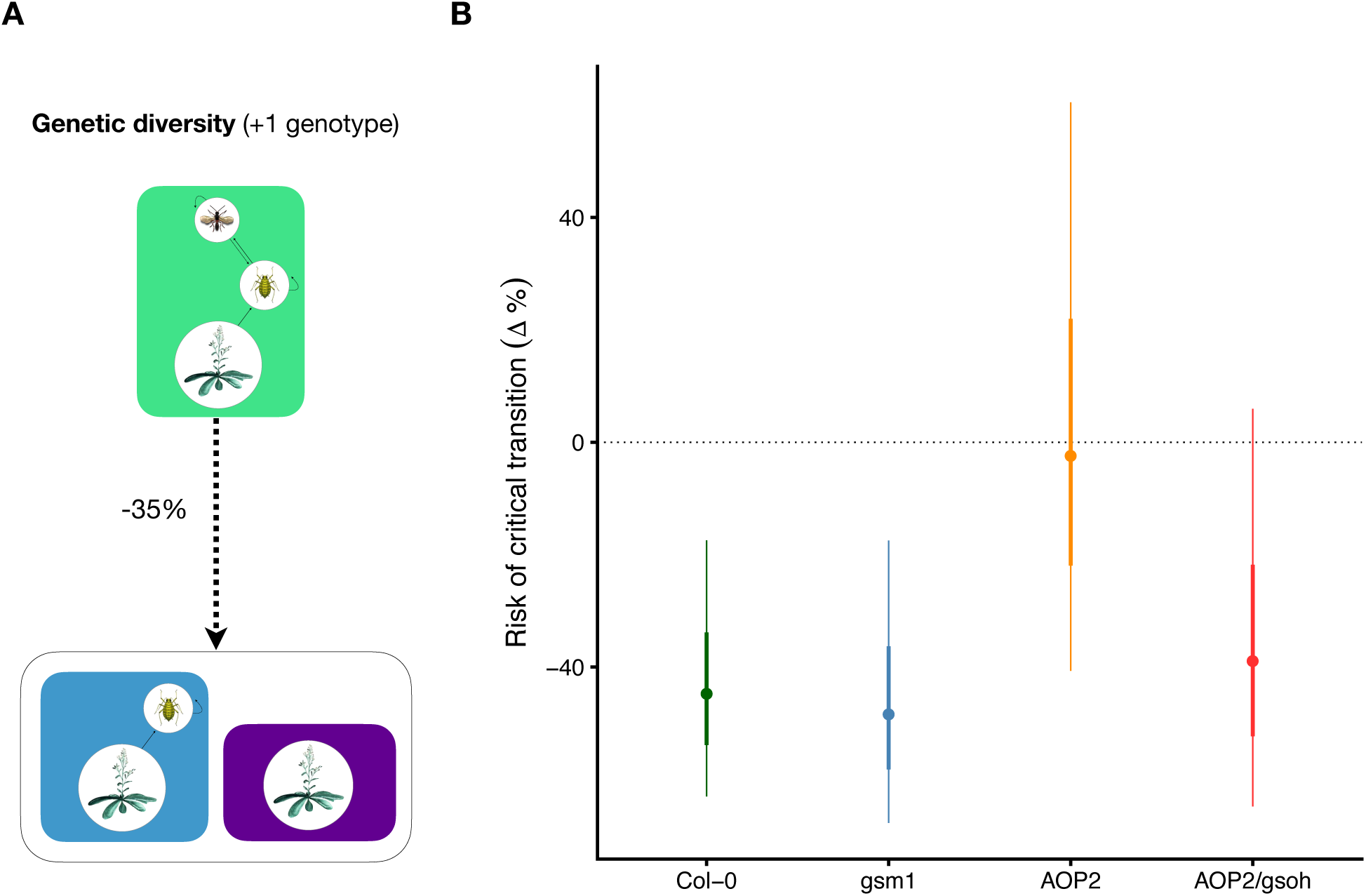
Identifying a keystone gene. (**A**) Genetic diversity reduced the overall risk of a critical transition in the food chain (green) to either an aphid only (blue) or completely collapsed (purple) state by 35%. (**B**) We then isolated the contribution of each genotype to this genetic diversity effect. Points correspond to mean estimates, while thick and thin bars correspond to standard errors and 95% confidence intervals, respectively. Note that the effect of genetic diversity (−35%) corresponds to the average genotype-specific effect (i.e., average of green, blue, red, and orange points). Both Col-0 and gsm1 have a non-functional *AOP2* gene and clearly decrease the risk of a critical transition. In contrast, both AOP2 and AOP2/gsoh have a functional *AOP2* gene and unclear effects. This suggests that *AOP2* functions as a keystone gene in this food web.

To understand how genetic change translates into food-web persistence, we applied theory on the structural stability of food webs. Structural stability quantifies the range of ecological conditions that allow species to stably coexist (*24, 25*). At the boundary of this range, the food web undergoes a critical transition to a simpler community (*21*). The geometry of intra- and interspecific interactions define the location of critical boundaries, while species’ intrinsic growth rates determine the proximity of the food web to a critical boundary. The distance, or to be precise the normalized angle, from a critical boundary measures the vulnerability of a food web to a critical transition. We used our time series data to quantify the effect of genetic diversity on interactions and species’ intrinsic growth rates, and thus the risk of a critical transition (details in Supplementary Material and Table S3).

We found that genetic diversity buffered the remaining food chain from a critical transition in a specific way (Fig. 4). This buffering effect was not because genetic diversity altered the location of the critical boundary (border of grey area in Fig. 4). Rather it was because genetic diversity moved the vector of intrinsic growth rates (solid arrows) upward and into the region where the parasitoid and aphid coexist (grey area in Fig. 4; Bayesian multivariate autoregressive (MAR(1)) model: 98% of posterior estimates > 0, Fig. S5). This buffering effect was primarily due to a concordant increase in the aphid’s intrinsic growth rate (Bayesian MAR(1) model: 97% of posterior estimates > 0), and to a lesser extent the parasitoid’s growth rate (Bayesian MAR(1) model: 87% of posterior estimates > 0). Although this analysis assumes that this food chain is at a stable equilibrium, our results hold under non-equilibrium conditions (i.e., persistence from different initial conditions, Fig. S6).

**Fig. 4.**
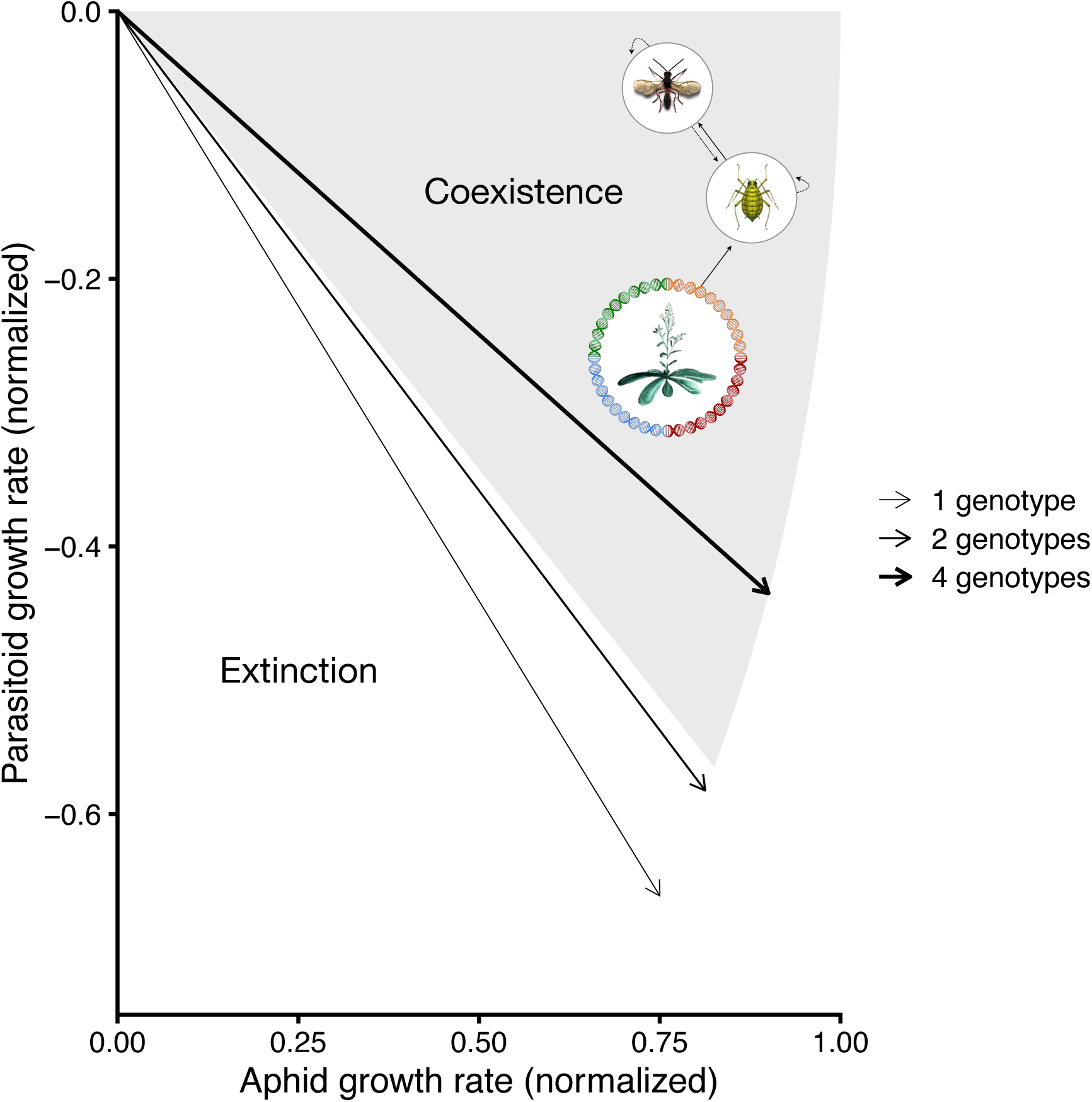
Genetic diversity buffers the remaining food chain from its critical boundary. The grey area represents the range of intrinsic growth rates (normalized to length 1) where the aphid and parasitoid coexist. The white area represents the range of intrinsic growth rates where the parasitoid goes extinct. Increasing genetic diversity moves the vector of intrinsic growth rates (solid arrows) upward and into the region of parameter space where the aphid and parasitoid coexist (grey area). Note that normalizing the vector of intrinsic growth rates to length 1 is necessary for accurately visualizing and calculating the proximity of the food chain to its critical boundary.

The buffering effect of genetic diversity was underlied by Col-0 and gsm1’s positive effect on aphid and parasitoid intrinsic growth rates (Fig. S7). This positive relationship was likely due to the faster relative growth rate of Col-0 and gsm1 genotypes compared to genotypes with a functional *AOP2* gene (Fig. S8; note that genetic diversity did not affect plant growth, ANOVA: *F*_1,9_ = 0.10, *P* = 0.759, Table S4). Work in similar environmental conditions has shown that natural accessions of *Arabidopsis* with a functional *AOP2* gene produce more total aliphatic glucosinolates and have slower growth rates (*10*), which has been shown to reduce the intrinsic growth rate of other aphid species (*26*). Note that the *AOP2* gene also has known pleiotropic effects on phenology, jasmonic acid signaling, and circadian rhythms (*27, 28*), which may ultimately mediate the dynamics of plant growth and aphid herbivory we observed. The direct plant genetic effect on the aphid’s growth rate is also the likely cause of the positive indirect effect on the parasitoid’s growth rate. For example, different plant genotypes of a closely related crucifer species alter the intrinsic growth rates of aphids and parasitoids in a coordinated manner (*29*). This positive correspondence in intrinsic growth rates is likely quite general among insect herbivores and their parasitoids (*30*).

These results hint at a general mechanism that may underlie a common pattern —plant populations with higher genetic diversity harbor more species-rich food webs (*31, 32*). Our work suggests that this pattern may be caused by an increased probability of having plants with chemical phenotypes that promote reproduction in herbivores and their natural enemies. While we have shown that this mechanism prevents critical transitions in a simple food chain, it is possible that this mechanism operates as well in more diverse food webs. Given that food chains of plants, insect herbivores, and their parasitoids comprise ∼40% of all described Eukaryotes (*33*), this mechanism may have far reaching consequences on the persistence and functioning of terrestrial ecosystems.

Taken together, our results show that genetic diversity in plant defense metabolism can increase the persistence of food webs in the face of climate warming. Although climate warming had a strong initial effect on food-web persistence, genetic diversity served as biological insurance against subsequent collapse. This suggests that the loss of genetic diversity we are witnessing (*34*) may accelerate the local extinction of species across multiple trophic levels. Yet, these results also present an opportunity for conservation and ecosystem restoration. For example, assisted migration of pre-adapted plant genotypes is becoming a well recognized strategy for restoring and preserving forest ecosystems into the future (*35*). Our results suggest that maximizing genetic diversity within pre-adapted populations may foster the structural integrity of terrestrial food webs in an uncertain and changing world.

## Acknowledgements

We thank Daniel Trujillo-Villegas and Xenia Muenger for their help conducting the experiment, and Matthias Furler for greenhouse support. Bernhard Schmid gave valuable feedback on the experimental design. Antonio Ferrera gave insight into quantifying the proximity of a critical boundary. This study was supported by the University Research Priority Program on Global Change and Biodiversity of the University of Zurich.

## Funding

Swiss National Science Foundation grant 31003A 160671 to J. Bascompte.

## Author Contributions

M.A. Barbour contributed to conceptualization, data curation, formal analysis, investigation, methodology, project administration, resources (insects), visualization, writing - original draft, writing - review and editing; D.J. Kliebenstein contributed to resources (plant genotypes), writing - review and editing; J. Bascompte contributed to conceptualization, funding acquisition, project administration, writing - review and editing.

## Competing interests

Authors declare no competing interests.

## Data and materials availability

All data and code to reproduce the reported results is publically available on GitHub (https://mabarbour.github.io/genes-to-foodweb-stability/) and have been archived on Zenodo (http://doi.org/10.5281/zenodo.3904601).

## Supplementary materials

Materials and Methods

Figs. S1 to S8

Tables S1 to S4

References *(36-58)*

## Notes

https://mabarbour.github.io/genes-to-foodweb-stability/

http://doi.org/10.5281/zenodo.3904601

### Competing Interest Statement

The authors have declared no competing interest.

