## Supplementary Material for "Genetic diversity increases food-web persistence in the face of climate warming"

### Supplementary materials for
### Genetic diversity increases food-web persistence in the face of climate warming

Matthew A. Barbour,[1*] Daniel J. Kliebenstein,[2] Jordi Bascompte[1]

[1]Department of Evolutionary Biology and Environmental Studies,
University of Zurich, 8057 Zurich, Switzerland
[2]Department of Plant Science, University of California Davis, Davis 95616 CA, USA

**This PDF file includes:**

Materials and Methods
Figs. S1 to S8
Tables S1 to S4

### Materials and Methods

#### Insect colonies

Prior to the experiment, we maintained colonies of the cabbage aphid *Brevicoryne brassicae*, the turnip aphid *Lipaphis erysimi*, and the parasitoid wasp *Diaeretiella rapae*, in a climate chamber at 20°C (16 h light/8 h dark photoperiod and 60% relative humidity). These insects were locally collected from *Brassica* spp. on the Irchel campus of the University of Zurich. Each aphid species was propogated from a single adult female and maintained continuous apomictic parthenogenesis under the abiotic conditions previously specified. Aphid populations were maintained on 2-week old seedlings of a non-experimental plant (radish, *Raphanus raphanistrum* subsp. *sativus* 'Saxa') in a circular pot (10D × 7.8H cm, 410 $cm^3$, GVZ Rossat) enclosed with a cellophane bag (160 × 300 mm, Celloclair). To ensure we had enough wingless adult aphids of similar age for the experiment, we distributed several adult aphids among several 2-week old radish seedlings (one adult aphid per seedling). After 48 h, we removed each adult, allowed nymphs to turn into adults 7–8 days later, and repeated this process for two additional weeks. The bulking population for each aphid was maintained in a large mesh cage (47.5W × 47.5D × 47.5H cm, BugDorm-4F4545 Insect Rearing Cage). The parasitoid population was maintained on a non-experimental aphid species (green peach aphid, *Myzus persicae*) that fed on the previously described radish variety in a large mesh cage. We also supplied parasitoids with multiple streaks of a honey solution (3:1 water:honey) at the top of the cage to promote survival and fecundity. Prior to the experiment, we split the parasitoid population into 2 separate populations (maintained under the same conditions) so we could add adult female parasitoids at consecutive weeks in the experiment. Note that we maintained aphids and parasitoids on non-experimental species to avoid any bias due to rearing conditions.

#### Plant genotypes

We used an existing set of transgenic lines (gsm1, AOP2, and AOP2/gsoh) that recreate natural knockout variation at three loci that control the biosynthesis of aliphatic glucosinolates in a Col-0 (natural accession) genetic background. The gsm1 genotype is an EMS mutant at the *MAM1* gene (At5g23010, ref. *13*). The AOP2 genotype is an insertion line (352:AOP2, ref. *36*) that gains a functional *AOP2* gene (At4g03060) and expresses *AOP2* to levels intermediate within natural *Arabidopsis* accessions (*14,18,37*). The AOP2/gsoh genotype is a manual cross between the AOP2 genotype and a T-DNA insertion line (SALK_09807, ref. *16*) that mimics a natural loss-of-function *GS-OH* allele (At2g25450, ref. *1*). Each of these transgenic lines has been backcrossed to Col-0 several times to remove unlinked polymorphisms in the original studies. The chemical phenotype produced by each of these genotypes is explained in Fig. 1B in the main text.

#### Experimental design

To manipulate genetic diversity, we created all possible genetic compositions ($n = 11$) of monocultures ($n = 4$), two ($n = 6$), and four genotypes ($n = 1$). We made 4 replicates for each monoculture, 6 replicates for each two-genotype mixture, and 8 replicates for the four-genotype mixture for a total of 60 experimental food webs. We could only fit 30 experimental food webs within a climate chamber, making this unbalanced design necessary to efficiently allocate experimental units among each genotypic composition.

To manipulate temperature for each experimental food web, we randomly split half of the replicates for each genetic composition between two climate chambers (Kälte 3000) set at different temperature regimes (20°C and 23°C), while other abiotic variables remained the same (16 h light/8 h dark photoperiod and 60% relative humidity). 20°C corresponds closely to the daily mean temperature these insects have experienced the past three years (2016–2019) in low elevation areas of northern Switzerland (<1,000 m) during their main activity period of June–August (19.3°C in ref. *38*). 23°C corresponds to the 3°C warming that northern Switzerland is predicted to experience in its daily mean temperature during the summer between 2045–2074 under current climate models (see Fig. 5 median JJA in ref. *19*). Therefore, these temperature treatments reflect the change in daily mean temperature this food web is expected to experience in the next 25–54 years. Note that these temperature regimes are not physiologically stressful for any of the species in these experimental food webs (*39–42*). To mitigate confouding effects of the temperature regime with the physical environment of the climate chamber, we exchanged experimental food webs between chambers once per month, but ensured that cages experienced the appropriate temperature regime.

To establish an experimental food web, we added 2 pots of *Arabidopsis* with the same genetic composition and total density (4 plants per pot) to a medium mesh cage (32.5W × 32.5D × 32.5H cm, BugDorm-4F3030 Insect Rearing Cage). We grew these plants in a nearby greenhouse chamber (mean daily mean 21.8°C, 18.5–24.7°C min.–max. daily mean) for three weeks prior to adding them to the experiment. To grow these plants, seeds of the appropriate genotype were sprinkled into one of four positions (randomly assigned) within a square pot (9L × 9W × 10H cm, GVZ Rossat) filled with damp soil (Einheitserde Classic). Pots were randomly distributed among eight slotted growing trays and covered with a plastic top for five days while seeds germinated. We thinned each pot once per week to maintain the appropriate genetic composition and total density. Pots were sub-irrigated twice per week for 15 minutes until they were moved into climate chambers, where they were watered once per week. Tray positions in the greenhouse were randomly shuffled twice per week (after watering) to avoid spatial variation in growing conditions.

Immediately after adding the 2 pots of *Arabidopsis* to a cage, we randomly distributed 2 wingless adult aphids (7–8 days old) of each species to each pot and randomly positioned the experimental food web in its appropriate temperature regime. Aphid populations were allowed to grow in the absence of parasitoids for 2 weeks, at which point we added 2 female parasitoids and repeated this at week 3. This ensured an overlap of parasitoid generations. Female parasitoids had been adults for 2–3 days when added to the experiment and presumably mated.

To ensure a constant resource supply to the aphids throughout the experiment, we replaced initial pots of *Arabidopsis* after two weeks, then once per week thereafter, with four week old plants of the same genetic composition and total density that were grown in the greenhouse (see paragraph before last for growing methods). We waited two weeks to replace the initial plants to minimize disturbance during initial aphid population growth. After counting the number of aphids and parasitoids (details in **Food-web dynamics**), we clipped and moved aboveground plant biomass to the back of the cage before replacing plants. This allowed aphids to disperse and find the new replacement plants. We kept plant biomass in the cage for 2 weeks to allow any parasitoids in mummified aphids to complete their development.

In a separate experiment, we assessed the direct effects of our experimental treatments on plant growth. We used the same procedure previously described for the initial pots, but we did not add insects to the cages. After two weeks in the climate chamber, we clipped aboveground plant biomass, oven-dried it at 60°C for 72 h, and weighed it to the nearest 1 mg.

#### Food-web dynamics

To quantify food-web dynamics, we surveyed aphid and parasitoid population abundances in each experimental food web every 7 days for 17 weeks. Plants were visually inspected and we counted the number of each aphid species, parasitized aphids (aphid mummies), and adult parasitoids. Due to the often high abundance of aphids, we counted aphids to a resolution of 5 individuals, whereas mummified aphids and parasitoids were counted to the individual level. Aphid mummies were not easily distinguished between species so these counts represent the total number of parasitized aphids. For statistical analysis, we combined counts of mummified aphids and adult parasitoids to estimate parasitoid abundance. To minimize disturbance, we did not count aphids and mummies underneath leaves of the basal rosette. This means that zero counts were not necessarily evidence of extinction. However, species that had zero counts for at least 3 weeks were classified as extinct and we used the first zero count as the time of extinction. For time points where species had zero counts, but were observed in subsequent weeks, we replaced zeros with the minimum resolution of our counts (i.e., 5 for aphids, and 1 for parasitoids).

To quantify variation in plant biomass over time, we collected biomass every 7 days, oven dried it at 60°C for 72 h, and weighed it to the nearest 1 mg. Note that collected biomass was taken from clipped plants at the back of the cage (details in **Experimental design**).

#### Statistical analyses of critical transitions, resulting food webs, and plant growth

To test the effect of genetic diversity, climate warming, and their interaction, on critical transitions in food-web structure, we conducted a discrete-time survival analysis using generalized linear models (GLM). A critical transition occured in our food web when a species went extinct, resulting in a simpler food web (see Fig. 2 in main text). Note that we frequently observed food webs with only the parasitoid in the week prior to complete collapse. Since the parasitoid invariably went extinct without its aphid resource, we classified these parasitoid only food webs as having completely collapsed when they were observed. The presence/absence of a critical transition to a simpler food web was modeled with a quasibinomial error distribution and a complementary log-log link function. The quasi-form of the binomial error distribution is the appropriate form for conducting $F$-tests (*43*), which we describe below. We used a complementary log-log function instead of the more familiar logit function as it has a more intuitive interpretation in survival analysis (*44*). That is, exponentiated coefficients can be interpreted as changes in the probability of an event, rather than the odds of the event. We specified each sampling interval as a discrete factor (`fweek`) and included plant genetic diversity (number of genotypes, `rich`), temperature (`temp`), and their statistical interaction (`temp:rich`) as predictor variables (Tables S1–S2). To test the effect of our experimental treatments, we used $F$-tests in a sequential (i.e., Type I) analysis of deviance (GLM ANOVA). Analysis of deviance is the extension of analysis of variance (ANOVA) for GLMs (*43*). This analysis allowed us to test the effect of our experimental treatments against their appropriate error terms and avoid pseudoreplication (*45*). Specifically, we tested the amount of deviance explained by `rich` compared to genetic composition (`com`), and `temp` and `temp:rich` compared to temperature by genetic composition (`temp:com`). Generalized linear mixed models (GLMMs) provide an alternative approach, but are often not appropriate for analyzing biodiversity experiments (*46*). This is because variance components (error terms previously mentioned) can be constrained to zero, resulting in tests of our experimental treatments against the residual deviance and degrees of freedom rather than their appropriate error term (i.e., pseudoreplicated tests). We observed this when we applied GLMMs to our data, which is why we chose to use GLM ANOVAs. To estimate uncertainty in the effect size (coefficients) of our experimental treatments, we calculated cluster-robust standard errors and 95% confidence intervals (*47*). We applied bias-reduced linearization to account for the relatively small number of clusters for each error term (11 `com` and 22 `temp:com`) and reduce the risk of Type I error (*48*). If we detected a clear effect of genetic diversity in a GLM ANOVA ($P < 0.05$), then we calculated the effect size of each specific genotype and its uncertainty with `com` as the cluster level. To do this, we replaced genetic diversity with four terms that indicate the presence/absence of each genotype. The average of these genotype-specific coefficients corresponds exactly to the coefficient for genetic diversity; therefore, this analysis allowed us to identify the contribution of each genotype to the observed effect of genetic diversity.

To test the effect of genetic diversity and climate warming on the resulting food web at the end of the experiment, we used a multinomial GLM and an analysis of deviance (GLM ANOVA) as previously described. We did not test the statistical interaction between plant genetic diversity and temperature (`temp:rich`) as one of the final food-web structures occurred less than 10 times (food chain = 7 cages; aphid only = 25 cages; complete collapse = 28 cages), making it unwise to test the `temp:rich` effect. Note, however, that `temp:rich` never had a clear effect on any of the critical transitions (all $P > 0.45$ in Tables S1–S2), suggesting our inference is robust.

To test the effect of genetic diversity, climate warming, and their interaction, on plant growth in the absence of insects, we applied a sequential ANOVA on log-transformed plant biomass (Table S4). As with the analysis of critical transitions, we calculated the effect of each specific genotype and its uncertainty using cluster-robust standard errors with `com` as the cluster level. Although we analyzed log-transformed plant biomass, we exponentiated our estimates to present effect sizes on the original scale (Fig. S8).

#### Bayesian multivariate autoregressive model

Critical transitions are determined by species' intrinsic growth rates and the strength of intra- and interspecific interactions (*21*). We used a multivariate autoregressive model (MAR(1), ref. *49*) to quantify the effect of our experimental treatments on interactions and intrinsic growth rates. For the 3 insect species in our food web, the MAR(1) model took the form:

$$\mathbf{N}_{it} = \mathbf{I}\mathbf{N}_{i,t-1} + (\mathbf{r} + \mathbf{A}\mathbf{N}_{i,t-1})(G_i + T_i) + \mathbf{p}P_{it} + \mathbf{c}C_i + \mathbf{E}_{it}, \quad (1)$$

where the elements of the model are as follows (vectors and matrices denoted in bold font):

- $\mathbf{N}_{it}$ is a $3 \times 1$ vector of population abundances (log transformed) in cage *i* at time *t*. We added a small positive constant (+1) prior to log transforming in order to include time points where species went extinct at time *t*.

- $\mathbf{I}$ is the $3 \times 3$ identity matrix. Ecologically, $\mathbf{I}\mathbf{N}_{i,t-1}$ accounts for the abundance of conspecifics at the previous time step. This effectively transforms our response variable into per-capita population growth rate. Note that we also added a small positive constant (+1) before log transforming $\mathbf{N}_{i,t-1}$ in order to maintain consistency with our response variable $\mathbf{N}_{i,t}$.

- $\mathbf{r}$ is a $3 \times 1$ vector of constants. Ecologically, $\mathbf{r}$ is a vector of intrinsic growth rates as it estimates per-capita population growth when other species and conspecifics are at low abundances (i.e., when $\mathbf{N}_{i,t-1} = 0$).

- $G_i$ is the genetic diversity of the plant population in cage *i*, where 0 = monoculture, 1 = two-genotype mixture, and 3 = four-genotype mixture.

- $T_i$ is the temperature regime of cage *i*, where 0 = 20°C and 3 = 23°C.

- $\mathbf{A}$ is a $3 \times 3$ matrix whose elements $a_{ij}$ give the effect of the log abundance of species *j* on per-capita population growth of species *i*. Ecologically, $\mathbf{A}$ is a matrix of intra- and interspecific interaction strengths.

- $P_{it}$ is plant biomass (log transformed) in cage *i* at time *t* and $\mathbf{p}$ is a $3 \times 1$ vector whose elements give the effect of plant biomass on per-capita population growth of each species. Ecologically, $P_{it}$ accounts for weekly variation in resource input into the food web.

- $C_i$ is the identity of cage *i* and $\mathbf{c}$ is a $3 \times 1$ vector whose elements give the effect of cage identity on per-capita population growth of each species. Statistically, $C_i$ functions as a random intercept term.

- $\mathbf{E}_{it}$ is a $3 \times 1$ vector of process errors for cage *i* at time *t* that has a multivariate normal distribution with mean vector $\mathbf{0}$ and covariance matrix $\sum$. Ecologically, $\mathbf{E}_{it}$ represents the effect of stochastic variation on the per-capita population growth of each species.

It does not make sense to try to estimate the effect of other species and conspecifics on the per-capita growth rate of an extinct species. Attempting to do so could severely bias parameter estimates. Therefore, we only fit the previously described MAR(1) model to data for which all three species were present in each cage at the previous time step (i.e., from week 2 until we detected the first critical transition at time *t* for cage *i*). This constraint, however, afforded us an opportunity to cross-validate our model. Specifically, we were able to test the predictive power of our model beyond the critical transition from the initial food web. The ability for our model to predict out-of-sample data guided our model selection (details in **Model selection**).

We analyzed our Bayesian MAR(1) models using the *brms* package in R version 3.6.2 (*50, 51*). This package uses the No-U-Turn Sampler for Markov Chain Monte Carlo (MCMC) sampling in the probabilistic programming language Stan (*52, 53*). Stan (via *brms*) provides a number of diagnostics for samples from the posterior distribution, including $\hat{R}$, effective sample size, measures of effective tree depth, and divergent transitions. We adjusted parameters controlling the maximum tree depth, the sampler's step size during the adaptation period, and the number of iterations until the model converged with no warnings. We found that running four chains for 4000 iterations each (discarding the first 2000 iterations as burn-in), specifying a step size of 0.8, and a maximum tree depth of 10, were sufficient for all models to converge with no warnings. For all parameters in the models we explored, $\hat{R}$ values were = 1.00, effective sample sizes were $>$750, and there were no divergent steps reported. A summary of the priors we used in our model is provided in Table S3. We also give a detailed justification of our choice of priors in the next section.

#### Choice of priors

Each of our prior estimates had a normal distribution $\mathcal{N}(\mu, \sigma)$, described by mean $\mu$ and standard deviation $\sigma$. We used life table studies of our aphid species to specify an informative prior for their intrinsic growth rates (*54, 55*). We specified $\sigma = 1$ to reflect our expectation that these intrinsic growth rates may differ considerably for our aphid populations on *Arabidopsis thaliana*. For the parasitoid, its intrinsic growth rate must be negative (i.e., in the absence of aphids, the parasitoid population will go extinct), which is reflected in our prior, $\mathcal{N}(-1.5, 1)$.

We used general ecological knowledge to choose priors for interaction ($a_{ij}$) and biomass effects ($p_i$). For example, we set a small negative effect ($\mu = -0.1$) for intraspecific interactions. This reflects our prior expectation of intraspecific competition rather than facilitation. Similarly, we set small negative aphid→aphid effects, and parasitoid→aphid effects. We set a small positive effect for each aphid→parasitoid effect and for plant biomass→aphid effects. Plant biomass may have positive or negative effects on parasitoids (*56*), so we set $\mu = 0$. For all interaction and biomass effects, we set $\sigma = 0.5$. This ensured that most of the prior distribution was between -1 and 1. This reflects our expectation that increasing densities would have a saturating effect on per-capita growth rate on an individual scale (e.g., Type 2 functional response).

It was unclear to us *a priori* which parameters would be affected by genetic diversity and temperature. Therefore, we set $\mu = 0$ and $\sigma = 0.5$ for each of these parameters. This type of prior makes the data "work" to show evidence for an effect. For the random effect of cage, we used a half-normal prior with $\mu$ informed by the standard deviation of aphid intrinsic growth rates from the life table studies (*54, 55*).

#### Model selection

Our goal was to identify a model that reproduced our observed effects of genetic diversity, specific genotypes, and temperature on critical transitions. Specifically, we sought a model where temperature accelerated a critical transition from the initial food web to a food chain without *B. brassicae* (as in Fig. 2 of main text). Simultaneously, we sought a model where genetic diversity, and more specifically Col-0 and gsm1, reduced the risk of a critical transition from the food chain with *L. erysimi* and *D. rapae* to either a complete collapse, or an aphid-only food web (as in Fig. 3 of main text). Remember that we only fit our model to data for which all three species were present (i.e., the initial food web); therefore, the effects of genetic diversity and specific genotypes were tested against data for which the model was not trained on (i.e., out-of-sample data). The full model described by equation 1 was not able to reproduce our observed effects of specific genotypes, as determined by the rank order effects of each genotype on the risk of a critical transition in the food chain. This suggests that the full model did not capture the underlying processes we observed.

To identify a better model, we removed parameters if 80% of their posterior estimates included zero, making sure to preserve marginality in higher-order terms (e.g., statistical interactions). This reduced model still could not reproduce the observed rank-order effects of specific genotypes on food-chain persistence. From here, we explored several alternative models that excluded some parameters, but not all, if 95% of their posterior estimates included zero. Finally, we also assessed a model for which all higher-order terms had 95% of their posterior estimates different from zero.

Following this procedure, we identified a single model that reproduced the observed effects of specific genotypes and genetic diversity (for out-of-sample data), as well as temperature (in-sample data). Note that there were other models that provided an equally good fit to the in-sample data, which described the critical transition from the initial food web to the food chain.

Only one model, however, was able to simultaneously reproduce our out-of-sample observations of genetic diversity and specific genotype effects on critical transitions in the food chain. This model took the following form:

$$\mathbf{N}_{it} = \mathbf{I}\mathbf{N}_{i,t-1} + \begin{bmatrix} r_B T_i \\ r_L G_i \\ r_D(G_i + T_i) \end{bmatrix} + \begin{bmatrix} \alpha_{BB} T_i & 0 & \alpha_{BD} T_i \\ \alpha_{LB} G_i & \alpha_{LL} & \alpha_{LD} \\ \alpha_{DB}(G_i + T_i) & \alpha_{DL} & 0 \end{bmatrix} \mathbf{N}_{i,t-1} + \begin{bmatrix} 0 \\ p_L \\ p_D \end{bmatrix} P_{it} + \mathbf{c} C_i + \mathbf{E}_{it}, \tag{2}$$

where subscripts *B*, *L*, and *D* indicate the aphid *B. brassicae*, the aphid *L. erysimi*, and the parasitoid *D. rapae*, respectively; zeros indicate coefficients that did not meet our previously described thresholds and were thus excluded from the model. We only expanded out matrices and vectors for which this model differed from the full model presented in equation 1. Note that when testing genotype-specific models, we replaced $G_i$ with Col + gsm1 + AOP2 + AOP2/gsoh, where each genotype = 1 if it was present in the plant population, and zero if absent.

### Structural stability analysis

Theory on the structural stability of ecological communities has been developed for deterministic models (*24, 25, 57, 58*). To leverage this theory, we translated our Bayesian MAR(1) model (equation 2) into a deterministic version. To do this, we ignored stochasticity ($\mathbf{E}_{it} = \mathbf{0}$) and condition our inferences on a common resource input (when oven-dried plant biomass = 1 g, $P_{it} = \ln(1) = 0$) for the average cage ($\mathbf{c} = \mathbf{0}$). Conditioning on different values may effectively shift the intrinsic growth rates, but will not alter our inferences about genetic diversity and temperature. If we then focus on a particular combination of experimental treatments (e.g., $G_i$ = 3 and $T_i$ = 0), our model reduces to:

$$\mathbf{N}_t = \mathbf{r} + (\mathbf{I} + \mathbf{A})\mathbf{N}_{t-1}. \tag{3}$$

From this model, we can estimate the vector of equilibrium abundances $\hat{\mathbf{N}}_\infty$ as:

$$\hat{\mathbf{N}}_{\infty} = -\mathbf{A}^{-1}\mathbf{r}. \tag{4}$$

Ecologically, $-\mathbf{A}^{-1}$ defines the range of intrinsic growth rates $\mathbf{r}$ that are compatible with species coexistence ($\hat{\mathbf{N}}_{\infty} > 0$). Moreover, we can identify the proximity of a food web to its critical boundary. To do this, we first describe the interaction matrix with column vector notation:

$$\mathbf{A} = \begin{bmatrix} \cdot & \cdot & \cdot \\ \mathbf{v}_B & \mathbf{v}_L & \mathbf{v}_P \\ \cdot & \cdot & \cdot \end{bmatrix}, \tag{5}$$

where subscripts *B*, *L*, and *D* indicate the aphid *B. brassicae*, the aphid *L. erysimi*, and the parasitoid *D. rapae*, respectively; each column vector $\mathbf{v}$ represents the effect of a species on itself and other species in the food web. The negative of each column vector defines a point, and the plane between two points defines a critical boundary of this food web (*58*). If we normalize each column vector to length one (e.g., $||\mathbf{v}_B|| = 1$), then we can visualize the critical boundaries of this food web on a sphere with a radius of 1 (as in Fig. S1).

Using *L. erysimi* and *D. rapae* as an example, the plane between $-||\mathbf{v}_L||$ and $-||\mathbf{v}_P||$ defines one of three critical boundaries in the initial food web. We then calculate their cross product to get the normal vector of this critical boundary, $||\mathbf{v}_{LP}|| = -||\mathbf{v}_L|| \times -||\mathbf{v}_P||$. Since the normal vector is perpendicular to the plane of the critical boundary, we subtract the angle between the vector of normalized intrinsic growth rates $||\mathbf{r}||$ and the normal vector of the critical boundary $||\mathbf{v}_{LP}||$ (through the origin $\mathbf{0}$) from $90°$. A normalized angle of zero indicates the food web lies on the critical boundary. More positive angles indicate the food web is able to tolerate larger perturbations (toward this critical boundary) without undergoing a critical transition. In contrast, more negative angles indicate the food web has crossed the critical boundary and at least one species will go extinct. With our Bayesian MAR(1) model, we calculated uncertainty in the normalized angle from a critical boundary using our posterior estimates of interactions and intrinsic growth rates. We constrained posterior estimates to biologically reasonable values on occasions where the estimates were unrealistic. This issue only arose for temperature effects on the initial food web, but never for genetic diversity effects on the food chain. For example, this occured for some estimates of *B. brassicae*'s intrinsic growth rate at 23°C (if estimate was negative, set to value of 0.1), *B. brassicae*'s effect on itself at 20°C (if estimate was positive, i.e., facilitation, set to value of 0), and *B. brassicae*'s effect on the parasitoid at 23°C (if estimate was negative, set to value of 0). Statistically, the 95% credible intervals of these particular parameters overlapped with zero, so an alternative would have been to simply set the parameter to zero. However, our approach of constraining them to biologically realistic values, rather than setting unrealistic estimates to zero, allowed for more uncertainty in our estimates (i.e., a more conservative inference).

Note that the normal vector calculation is necessary for determining the proximity to a critical boundary for a food web with 3 or more species. If there are only two species, then the critical boundary reduces to a point. For example, in the persistent food chain, the critical boundary is determined by $-\|\mathbf{v}_L\|$. We then calculate the angle between $\|\mathbf{r}\|$ and $-\|\mathbf{v}_L\|$ (through the origin $\mathbf{0}$) to get the normalized angle from the critical boundary.

#### Non-equilibrium simulation

The structural stability analysis assumes the food web is at a locally stable equilibrium. Although useful, it is important to check that our inference holds under non-equilibrium conditions. This is especially true for our experiment given our frequent observations of critical transitions (i.e., non-equilibrium conditions).

We used the mean parameter estimates from our Bayesian MAR(1) model to simulate the population dynamics of each species. The initial abundance of species in the food web were experimentally controlled; therefore, we only used this one set of initial conditions to simulate the effects of our experimental treatments on the first critical transition in the initial food web. For the remaining food chain, we simulated a range of initial conditions based on observed abundances of *L. erysimi* and *D. rapae* after *B. brassicae* went extinct. We simulated food chain dynamics for 10 time steps, since this corresponded closely to the average number of weeks remaining in the experiment after *B. brassicae* went extinct. We set an extinction threshold so that if a species' log abundance was less than 1 (i.e., 2–3 individuals), then it was classified as extinct.

### Figs. S1 to S8

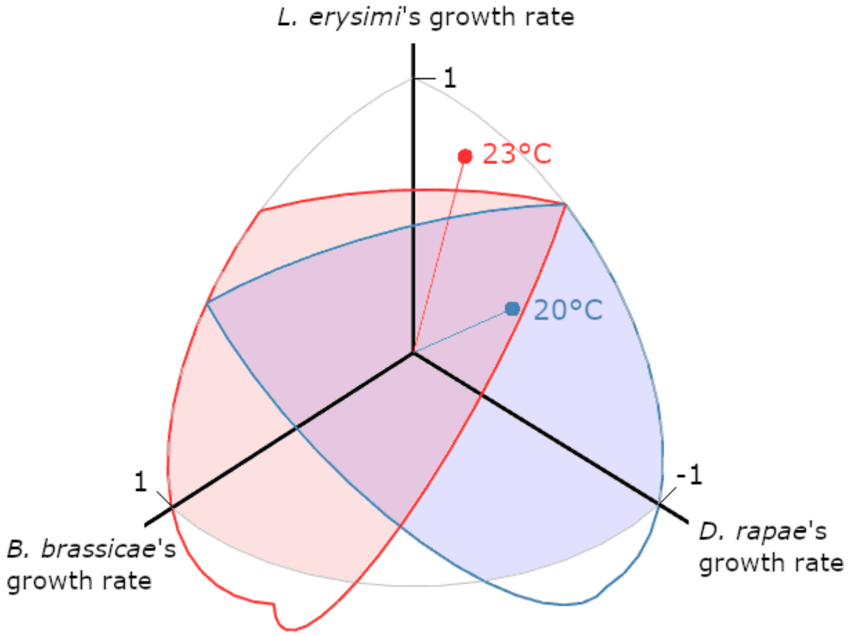


**Fig. S1. Warming increased the risk of a critical transition in the initial food web.** Each axis corresponds to the intrinsic growth rate of a species in the initial food web (normalized to a length of 1). We rotated the axes to focus on the range of realistic growth rates, i.e., positive growth rates for aphids, and a negative growth rate for the parasitoid. This range of realistic growth rates is circumscribed by the grey spherical triangle. The blue and red spherical triangles indicate the range of growth rates in which all species stably coexist at 20°C and 23°C, respectively. Blue and red points correspond to the vector of normalized intrinsic growth rates for all three species at 20°C and 23°C, respectively. Note that the growth rate vector of species at 23°C is outside the region where all species stably coexist (red-shaded region). This resulted in a more rapid extinction of *B. brassicae* in our warming treatment (Fig. S3). Once a species goes extinct, the range of growth rates in which the remaining two species coexist can be represented in 2-dimensions, as in Fig. 4 of the main text.

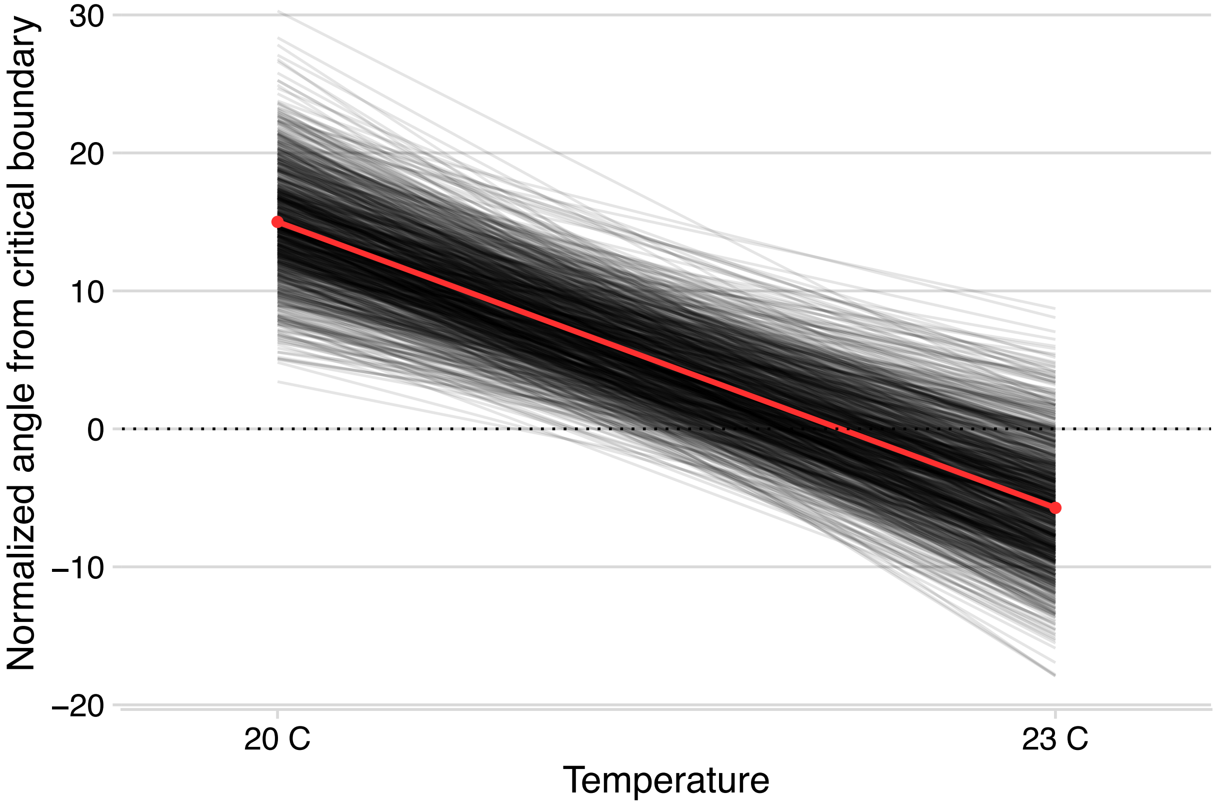


**Fig. S2. Proximity of the initial food web to its nearest critical boundary.** Each line corresponds to an estimate from the posterior distribution of our Bayesian MAR(1) model (equation 2), and the red line corresponds to the average effect. For visualization, we only plot 1000 (of 8000) posterior estimates, but our inference is based on all posterior estimates. More positive angles from the critical boundary reduce the risk of a critical transition, whereas negative angles increase the risk. Warming increased the risk of a critical transition from the initial food web toward a simpler food chain in 100% of posterior estimates.

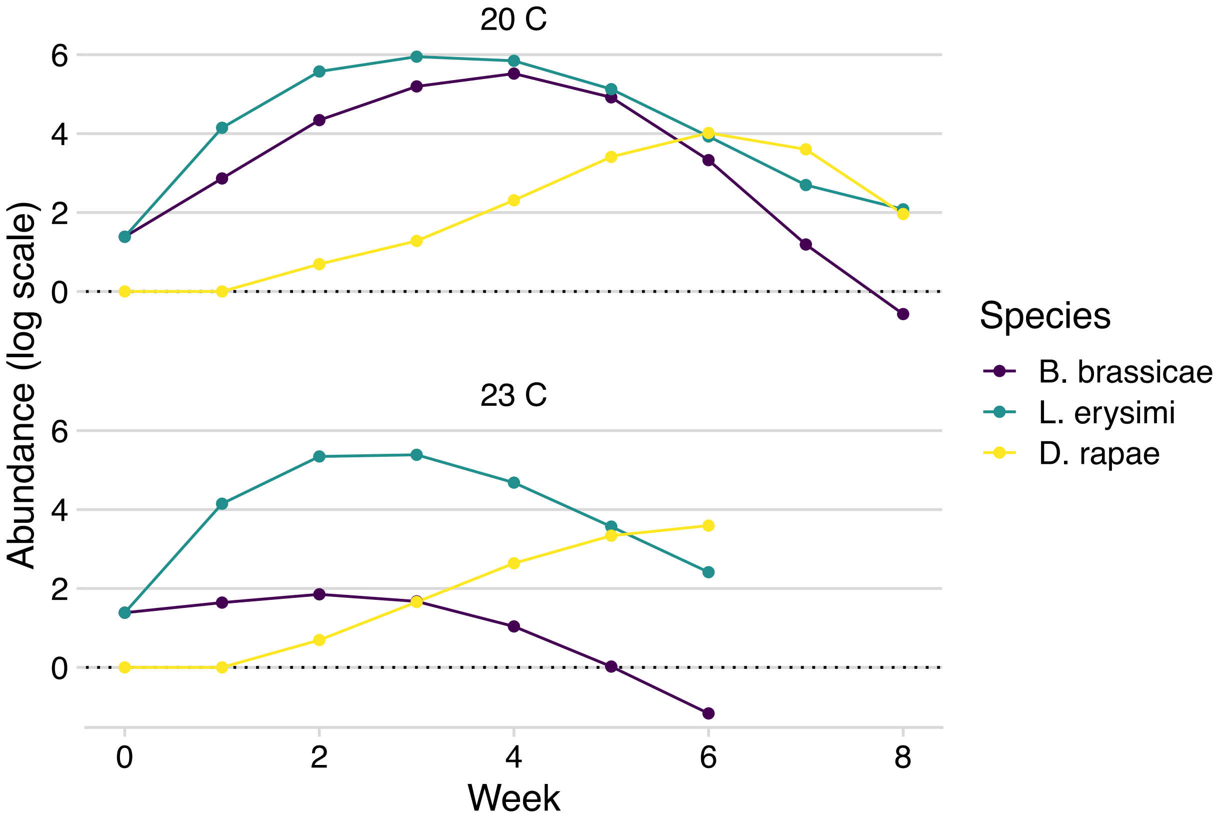


**Fig. S3. Initial food-web persistence under non-equilibrium conditions**. We found that the aphid *B. brassicae* went extinct in both temperature treatments, but this extinction occurs more quickly at 23°C. This corresponds to our observation that warming increased the risk of a critical transition from the initial food web toward a simpler food chain (see Fig. 2 in main text).

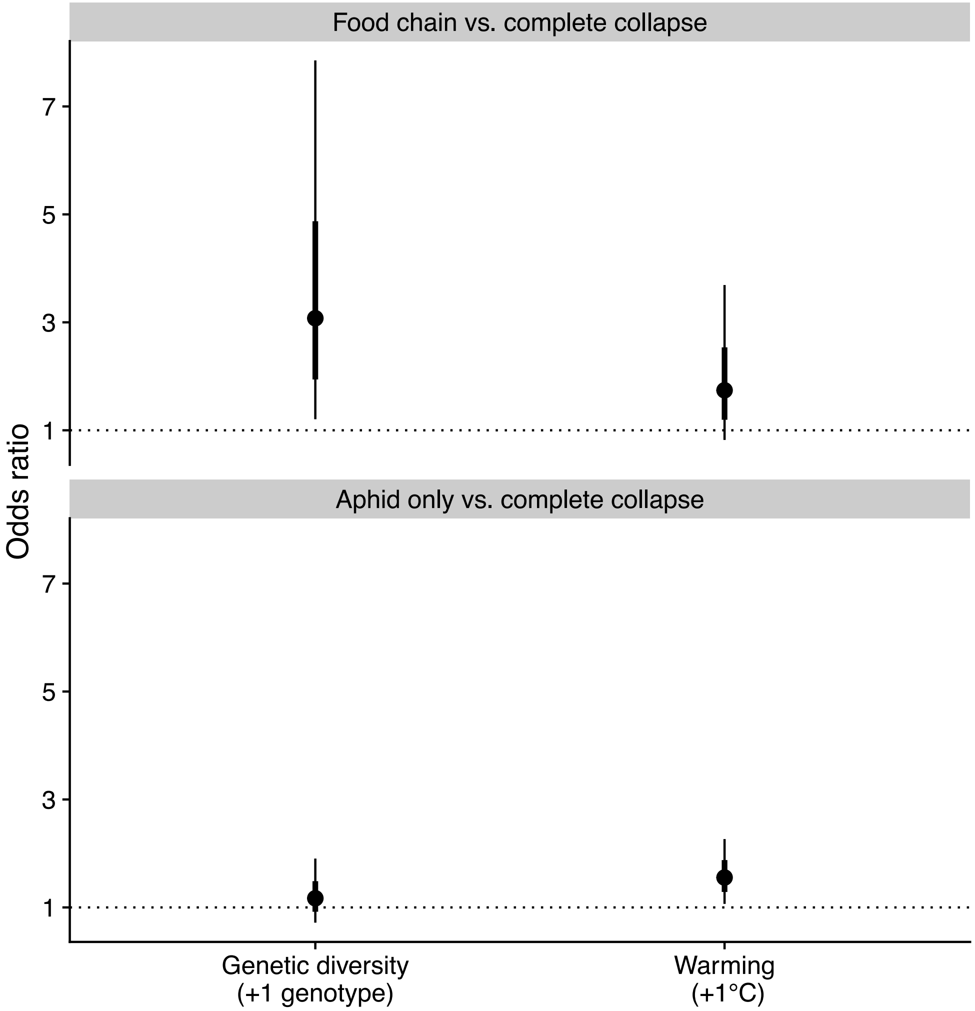


**Fig. S4. Odds of the different food webs resulting at the end of the experiment.** An odds ratio of 1 indicates no effect, whereas values greater (less) than 1 indicate an increase (decrease) in the odds of observing a food chain (top panel) or one aphid only (bottom panel) relative to complete collapse (i.e., all insects went extinct). Points correspond to mean estimates, while thick and thin bars correspond to standard errors and 95% confidence intervals, respectively.

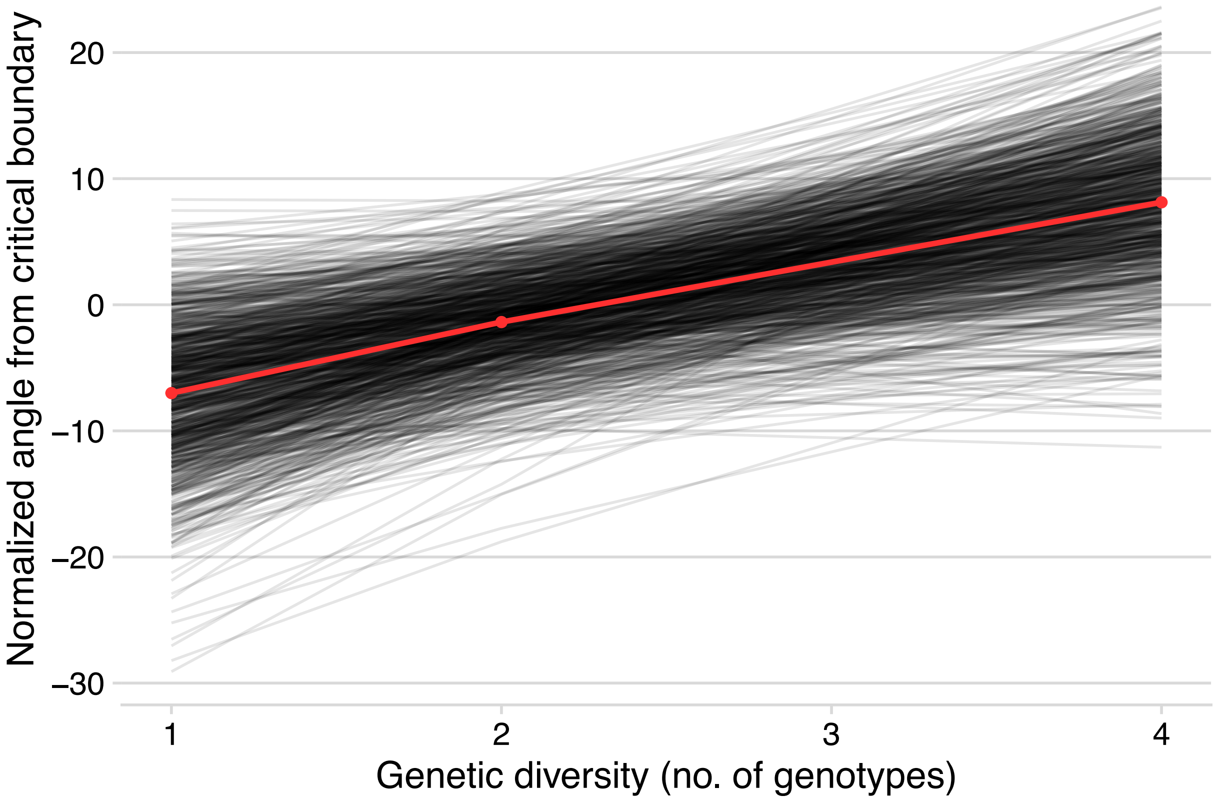


**Fig. S5. Proximity of the remaining food chain to its critical boundary.** Each black line corresponds to an estimate from the posterior distribution of our Bayesian MAR(1) model (equation 2), and the red line corresponds to the average effect. For visualization, we only plot 1000 (of 8000) posterior estimates, but our inference is based on all posterior estimates. More positive angles from the critical boundary reduce the risk of a critical transition, whereas negative angles increase the risk. Genetic diversity reduced the risk of a critical transition in 98% of posterior estimates.

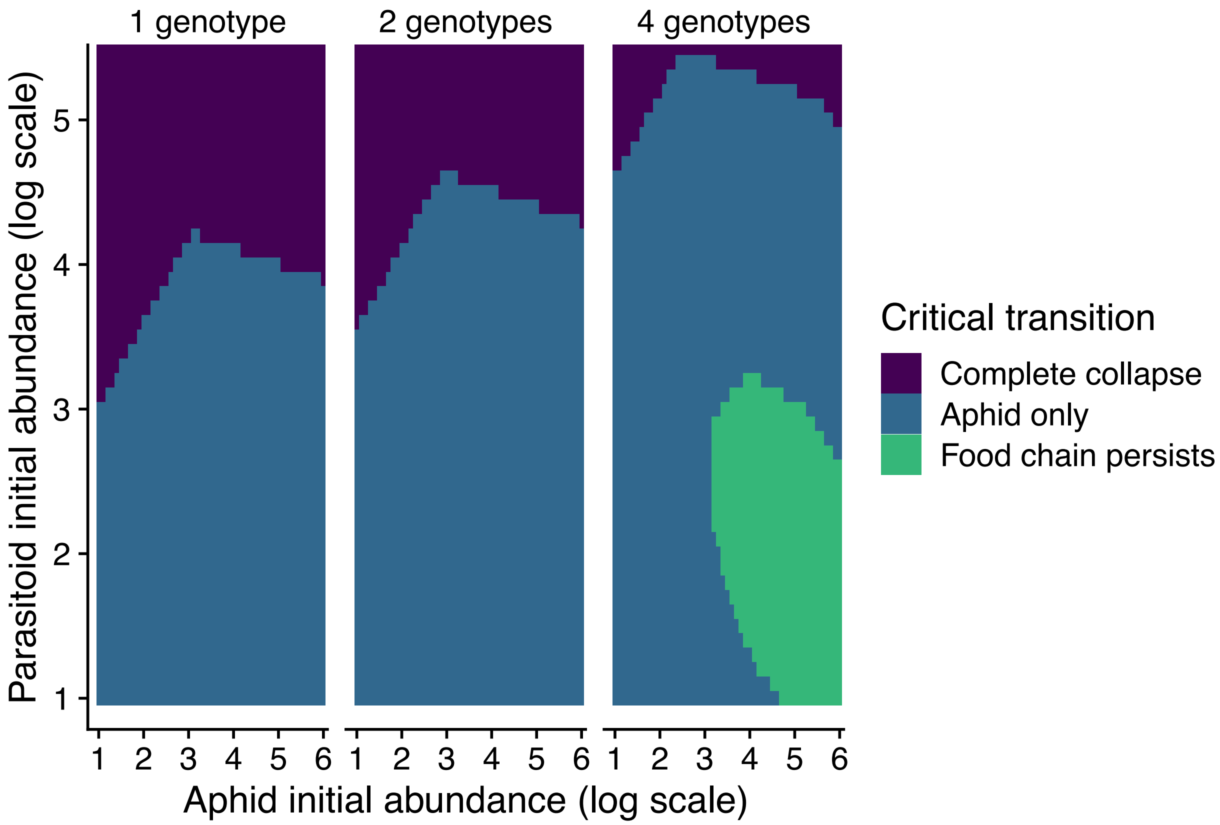


**Fig. S6. Food-chain persistence under non-equilibrium conditions**. We found that genetic diversity decreased the risk of both the aphid and parasitoid going extinct (complete collapse, purple region) and increased the likelihood that the food chain would persist (green region). This matches our observation that genetic diversity increased the odds of the food chain persisting relative to complete collapse (Fig. S4).

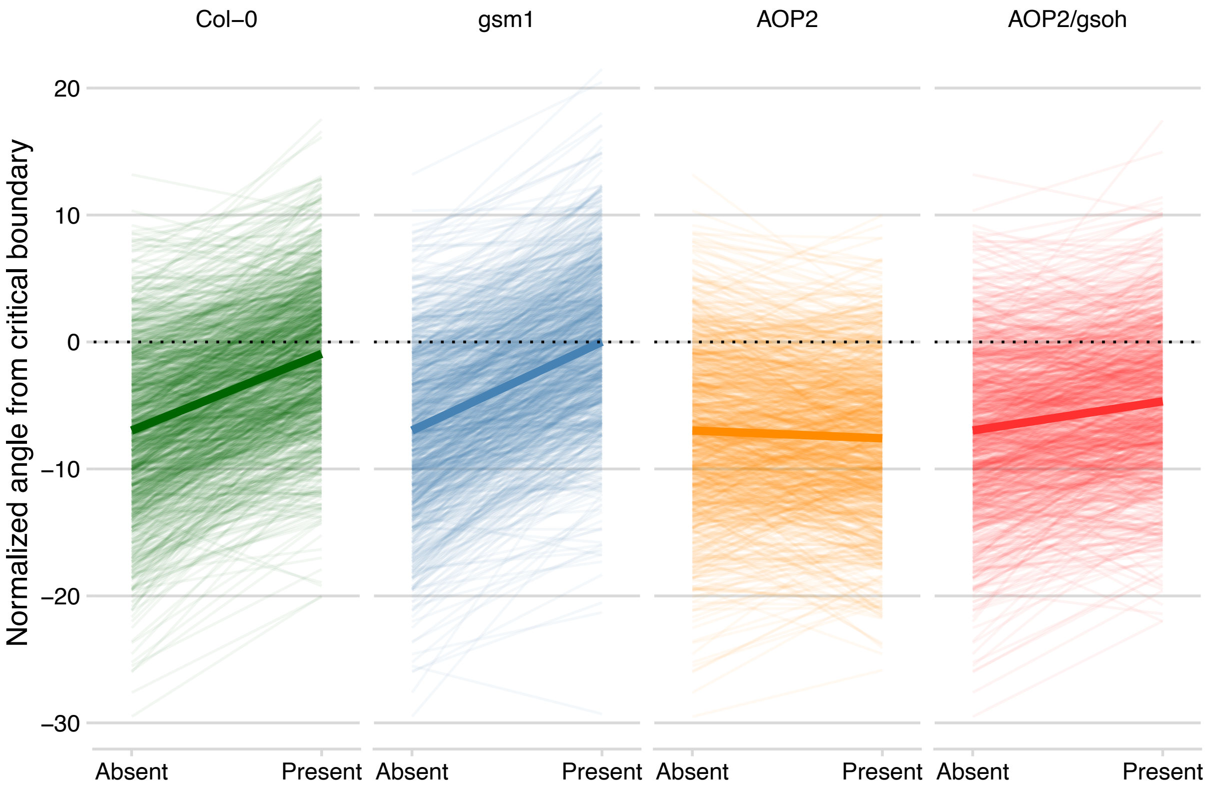


**Fig. S7. Plant genotype alters the proximity of the remaining food chain to its critical boundary.** Thin lines correspond to estimates from the posterior distribution of our Bayesian MAR(1) model (equation 2, but with genotypes substituted for $G_i$), whereas the thick line corresponds to the average effect. For visualization, we only plot 1000 (of 8000) posterior estimates, but our inference is based on all posterior estimates. More positive angles from the critical boundary reduce the risk of a critical transition, whereas negative angles increase the risk. Col-0 and gsm1 reduced the risk of a critical transition in 92% and 94% of posterior estimates, respectively. In contrast, AOP2 and AOP2/gsoh reduced the risk of a critical transition in only 43% and 70% of posterior estimates, respectively. Note that this pattern matches the observed effects of each genotype on the risk of a critical transition in the remaining food chain (see Fig. 3B in main text).

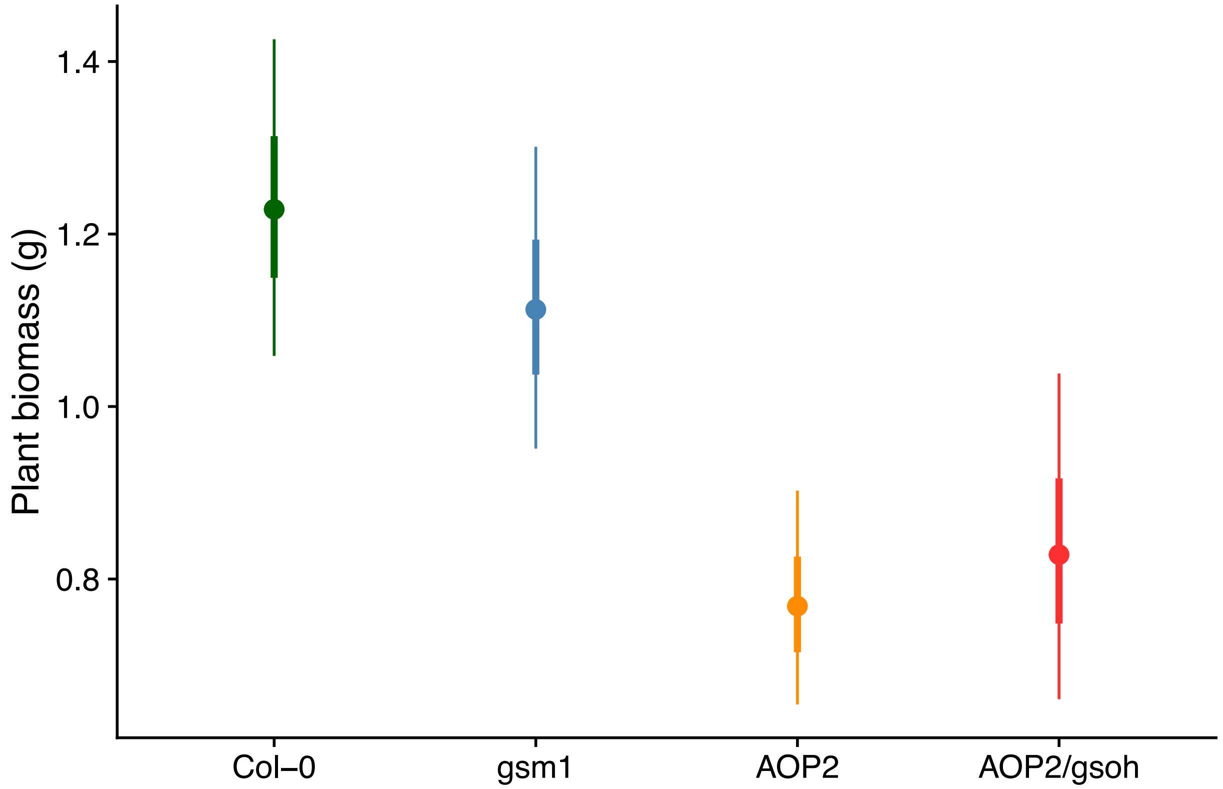


**Fig. S8. Genotype-specific effects on plant growth in the absence of insects.** Points correspond to mean estimates, while thick and thin bars correspond to standard errors and 95% confidence intervals, respectively. These estimates control for variation in temperature, but we plot them at the baseline temperature treatment of 20°C. While there was a clear positive effect of having a non-functional (Col-0 and gsm1) vs. functional (AOP2 and AOP2/gsoh) *AOP2* gene on plant growth, genetic diversity did not alter plant growth (Table S4).

### Tables S1 to S4

**Table S1.** Analysis of deviance for discrete-time survival GLM of critical transitions in the initial food web (yellow in Fig. 2).

| Source | df (Source) | df (Error) | Deviance | Mean Deviance | *F* | *P* | Error |
|---|---|---|---|---|---|---|---|
| Any critical transition<br>(yellow to either green or purple in Fig. 2) | | | | | | | |
| fweek | 4 | 177 | 65.23 | 16.31 | 20.854 | <0.001 | Residuals |
| temp | 1 | 9 | 14.81 | 14.81 | 11.132 | 0.009 | temp:com |
| rich | 1 | 9 | 2.18 | 2.18 | 1.713 | 0.223 | com |
| temp:rich | 1 | 9 | 0.66 | 0.66 | 0.498 | 0.498 | temp:com |
| Critical transition to the food chain<br>(yellow to green in Fig. 2) | | | | | | | |
| fweek | 4 | 177 | 35.83 | 8.96 | 10.916 | <0.001 | Residuals |
| temp | 1 | 9 | 20.19 | 20.19 | 11.202 | 0.009 | temp:com |
| rich | 1 | 9 | 0.38 | 0.38 | 0.407 | 0.539 | com |
| temp:rich | 1 | 9 | 0.38 | 0.38 | 0.210 | 0.658 | temp:com |
| Critical transition to complete collapse<br>(yellow to purple in Fig. 2) | | | | | | | |
| fweek | 1 | 21 | 0.64 | 0.64 | 0.631 | 0.436 | Residuals |
| temp | 1 | 6 | 0.93 | 0.93 | 1.056 | 0.344 | temp:com |
| rich | 1 | 9 | 0.42 | 0.42 | 0.262 | 0.621 | com |

**Table S2.** Analysis of deviance for discrete-time survival GLM of a critical transition from the remaining food chain (green in Fig. 2 & 3A in main text).

| Source | df (Source) | df (Error) | Deviance | Mean Deviance | $F$ | $P$ | Error |
|---|---|---|---|---|---|---|---|
| Any critical transition (green to either blue or purple in Fig. 3A) | | | | | | | |
| fweek | 9 | 252 | 18.05 | 2.01 | 2.544 | 0.008 | Residuals |
| temp | 1 | 9 | 0.01 | 0.01 | 0.005 | 0.946 | temp:com |
| rich | 1 | 9 | 5.32 | 5.32 | 9.006 | 0.015 | com |
| temp:rich | 1 | 9 | 0.85 | 0.85 | 0.592 | 0.461 | temp:com |
| Critical transition to aphid only (green to blue in Fig. 2) | | | | | | | |
| fweek | 9 | 252 | 9.99 | 1.11 | 2.094 | 0.031 | Residuals |
| temp | 1 | 9 | 0.36 | 0.36 | 0.238 | 0.637 | temp:com |
| rich | 1 | 9 | 0.81 | 0.81 | 0.508 | 0.494 | com |
| temp:rich | 1 | 9 | 1.07 | 1.07 | 0.715 | 0.420 | temp:com |
| Critical transition to complete collapse (green to purple in Fig. 2) | | | | | | | |
| fweek | 3 | 94 | 1.03 | 0.34 | 0.469 | 0.705 | Residuals |
| temp | 1 | 9 | 0.51 | 0.51 | 0.373 | 0.557 | temp:com |
| rich | 1 | 9 | 5.42 | 5.42 | 5.477 | 0.044 | com |
| temp:rich | 1 | 9 | 0.70 | 0.70 | 0.518 | 0.490 | temp:com |

**Table S3.** Summary of prior distributions.

| Prior distribution | Model term | Justification |
|---|---|---|
| $\mathcal{N}(\mu = 2.26, \sigma = 1)$ | $r$ for aphid *B. brassicae* | ref. *54* |
| $\mathcal{N}(\mu = 2.20, \sigma = 1)$ | $r$ for aphid *L. erysimi* | ref. *55* |
| $\mathcal{N}(\mu = -1.5, \sigma = 1)$ | $r$ for parasitoid *D. rapae* | Negative values realistic; assumed similar absolute value as aphid. |
| $\mathcal{N}(\mu = -0.1, \sigma = 0.5)$ | Intraspecific effect | Negative values realistic; set weak prior to regularize |
| $\mathcal{N}(\mu = -0.1, \sigma = 0.5)$ | Aphid $i \rightarrow$ aphid $j$ | Negative values realistic; set weak prior to regularize |
| $\mathcal{N}(\mu = -0.1, \sigma = 0.5)$ | Parasitoid $\rightarrow$ aphid $i$ | Negative values realistic; set weak prior to regularize |
| $\mathcal{N}(\mu = 0.1, \sigma = 0.5)$ | Aphid $i \rightarrow$ parasitoid | Positive values realistic; set weak prior to regularize |
| $\mathcal{N}(\mu = 0.1, \sigma = 0.5)$ | Plant biomass $\rightarrow$ aphid $i$ | Positive values realistic; set weak prior to regularize |
| $\mathcal{N}(\mu = 0, \sigma = 0.5)$ | Plant biomass $\rightarrow$ parasitoid | Regularizing toward zero; no prior expectation on direction |
| $\mathcal{N}(\mu = 0, \sigma = 0.5)$ | Genetic diversity effects | Regularizing toward zero; no prior expectation on direction |
| $\mathcal{N}(\mu = 0, \sigma = 0.5)$ | Temperature effects | Regularizing toward zero; no prior expectation on direction |
| half-$\mathcal{N}(\mu = 0.03, \sigma = 0.5)$ | $\sigma$ of Cage intercept | $\sigma$ in $r$ estimates from refs. *54, 55* |

**Table S4.** Analysis of variance for plant biomass (log transformed) in the absence of insects.

| Source | df (Source) | df (Error) | Deviance | Mean Deviance | *F* | *P* | Error |
|---|---|---|---|---|---|---|---|
| temp | 1 | 9 | 6.00 | 6.00 | 65.429 | <0.001 | temp:com |
| rich | 1 | 9 | 0.04 | 0.04 | 0.100 | 0.759 | com |
| temp:rich | 1 | 9 | 0.03 | 0.03 | 0.291 | 0.603 | temp:com |
